# Trimitomics: An efficient pipeline for mitochondrial assembly from transcriptomic reads in non-model species

**DOI:** 10.1101/413138

**Authors:** Bruna Plese, Maria Eleonora Rossi, Nathan James Kenny, Sergi Taboada, Vasiliki Koutsouveli, Ana Riesgo

**Affiliations:** Life Sciences Department, The Natural History Museum, Cromwell Road, London SW7 5BD, UK; Division of Molecular Biology, Ruder Bošković Institute, Bijenička cesta 54, 10000, Zagreb, Croatia

**Keywords:** Mitochondrial genome, transcriptomics, assembly, invertebrates

## Abstract

Mitochondrial resources are of known utility to many fields of phylogenetic, population and molecular biology. Their combination of faster and slower-evolving regions and high copy number enables them to be used in many situations where other loci are unsuitable, with degraded samples and after recent speciation events.

The advent of next-generation sequencing technologies (and notably the Illumina platform) has led to an explosion in the number of samples that can be studied at transcriptomic level, at relatively low cost. Here we describe a robust pipeline for the recovery of mitochondrial genomes from these RNA-seq resources. This pipeline can be used on sequencing of a variety of depths, and reliably recovers the protein coding and ribosomal gene complements of mitochondria from almost any transcriptomic sequencing experiment. The complete sequence of the mitochondrial genome can also be recovered when sequencing is performed in sufficient depth. We evidence the efficacy of our pipeline using data from a number of non-model invertebrates of four disparate phyla, namely Porifera, Nemertea, Mollusca and Annelida. Interestingly, among our poriferan data, where microbiological symbionts are known empirically to make mitochondrial assembly difficult, this pipeline proved especially useful.

Our pipeline will allow the recovery of mitochondrial data from a variety of previously-sequenced samples, and add an additional angle of enquiry to future RNA-seq efforts, simplifying the process of mitochondrial genome assembly for even the most recalcitrant clades and adding this data to the scientific record for a range of future uses.

## Introduction

Mitochondria, due to their vital function within the cell, have been the subject of research from a diversity of angles for as long as we have known of their existence. In particular, the invariant protein coding cassette, combination of fast and slow-evolving characters and severe functional constraints on their operation (da Fonesca et al., 2008), render them excellent for molecular phylogenetic investigations. Mitochondrial sequence data are exceptionally useful for tracing the inter-relationships of non-model organisms. They can also, among other examples, be used to understand population structure (Avise et al., 1987), for conservation biology (Rubinoff, 2006), for forensics (Melton et al., 2012), in medicine (Picard et al., 2016), and in the study of evolutionary pressures (Romero et al., 2016). As a result of this flexibility, the sequencing of mitochondrial genomes is popular for a diverse range of uses.

While there are a number of means by which mitochondrial genomes can be sequenced, including Sanger and genomic DNA sequencing, it has been noted previously that RNA-seq data could provide a novel source of mitochondrial sequence data (Smith, 2013; Tian & Smith, 2016). Despite treatments aimed at enriching nuclear genome mRNA levels, the high copy number of mitochondria within cells, coupled to generally high expression levels, mean that some level of mitochondrial data will inevitably be present in any transcriptomic dataset (Raz et al., 2011; Smith, 2013; Tian & Smith, 2016). This can be leveraged to extract useful sequence information from transcriptomic assembly (Tian & Smith, 2016). RNA-seq has been applied to almost every major clade in the tree of life, and has proven its utility repeatedly for solving myriad questions in life’s evolution. The extraction of mRNA from mixed or single tissues, followed by conversion to cDNA and the construction of libraries from this for sequencing, has become a standard technique in the study of many realms of molecular biology.

The basic technique of RNA-seq is often supplemented by a step which removes ribosomal and transfer RNA from the RNA sample, leaving mRNA in higher relative abundance. These poly-A enrichment techniques make use of the polyadenylation of RNA polymerase 2 products in eukaryotic cells (Hirose & Manley, 1998), although it is important to note that some prokaryote mRNAs are also polyadenylated (Régnier & Marujo, 2013). Polyadenylated mRNAs are bound to ligands (often with a poly-T “bait”), and subsequently separated from the remaining rRNA and tRNA fraction, which can comprise around 95% of the RNA in a cell. Mitochondrial RNA is polyadenylated in some branches of the tree of life but not others (Chang & Tong, 2012; Bratic et al., 2016), and this pattern is still not fully understood. In some clades, a polyadenylation signal marks RNA for degradation (Chang & Tong, 2012). Mitochondrial RNA will therefore be variably recovered after poly-A enrichment methods, depending on the clade from which the RNA was extracted, and the role of polyadenylation within that group. Other methods of rRNA removal, for instance “Ribozero” approaches, are also used, although less frequently.

A plethora of mitochondrial assembly tools are available, but these generally rely on gDNA or Sanger-derived reads, rather than those sourced from RNA-seq experiments. Programmes such as Norgal (Al-Nakeeb et al., 2017), MitoBIM (Hahn et al., 2013), and NOVOPlasty (Dierckxsens et al., 2016) have proven useful at recovering mitogenome assemblies, particularly when the sequence of a closely related individual is available for use as a map for assembly. Increasingly sophisticated tools are available for using gDNA for assembly (e.g. Schomaker-Bastos & Prosdocimi, 2008) However, previous approaches, and particularly those reliant on mining the results from transcriptomic assembly programs, are not always reliable when using mRNA sequence data as the basis for assembly, especially when using reads that have been subject to rRNA removal or mRNA enrichment (Tian & Smith, 2016).

Here we describe a novel pipeline for the reconstruction of mitochondrial gene cassettes and whole coding sequences from RNAseq reads, based on existing, freely available programmes. This pipeline uses a sequential approach and established tools, so that RNA-seq reads of good quality and uniform coverage across the mitogenome will quickly be assembled into workable data, while those datasets posing problems can still yield results, albeit with additional effort. We have benchmarked this process using reads derived from a number of Illumina platforms, read lengths and sequencing depths, and show that reliable mitochondrial genome assemblies, and particularly the sequence of protein coding genes, can be reliably recovered from even polyA selected samples. This method will allow the recovery of mitochondrial genome data from both historical and novel RNA-seq experiments, and provide a variety of novel resources to the scientific community for use in the myriad of applications for which mitogenomes have proven their utility.

## Materials and Methods

In this study we used RNA-seq data from representative species encompassing four phyla. We analysed four previously published RNA-seq datasets (two sponges, a mollusc, and an annelid) and also generated new data for one sponge and one nemertean. RNA-seq datasets for *Corticium candelabrum* and *Ircinia fasciculata* (Porifera), *Biomphalaria glabrata* (Mollusca) and *Platynereis dumerilii* (Annelida) were obtained from the NCBI Sequence Read Archive (SRA) under accession numbers SRR504694, SRP037543, SRR3039143, and SRR1742987, respectively. The species *C. candelabrum, B. glabrata*, and *P. dumerilii* already have published mitochondrial genomes (mt genomes), and these were used to estimate the accuracy and efficiency of the proposed workflow.

For the nemertean *Antarctonemertes valida* and the poriferan *Spongia officinalis*, new RNA-seq data were generated, and deposited in the SRA under the following accession numbers: SRP157324 and SRP150632. The newly assembled mt genomes for these two species, and that of *I. fasciculata,* have been submitted to GenBank under accession numbers MH768970-MH768972.

### RNA extraction, library preparation, Illumina sequencing

For *S. officinalis* (Koutsouveli et al., unpublished data), total RNA was extracted from 20 tissue pieces (4 different individuals and 5 replicates per individual) with TRIzol (Ambion) and mRNA purified with Ribo-Zero™ (Illumina). Library preparation of all 20 samples was performed with TruSeq Stranded Total RNA Library Prep Kit (Illumina) and further sequenced with an Illumina HiSeq 2000 using a paired-end (150 bp length) sequencing strategy.

A single individual divided into three portions (anterior part, posterior part, and proboscis) was used for *A. valida.* Total RNA extraction was performed using TRIzol (Ambion) and mRNA purification with Dynabeads mRNA DIRECT Purification kit (ThermoFisher Scientific) following already established protocols (see Riesgo et al., 2014; Kenny et al., 2018). Library preparation for *A. valida* was performed with the commercial ScriptSeq v2 RNA-seq library preparation kit (Epicentre) and the three libraries were sequenced on an Illumina NextSeq500 to a length of 150 bp (paired-end).

Details on the RNA extraction, library preparation and sequencing for *C. candelabrum* and *I. fasciculata* can be found in Riesgo et al., 2014, while those for *B. glabrata* are detailed in Kenny et al., 2016 and for *P. dumerilii* in Achim at al., 2018.

### Recovering mt genomes

The quality assessment of all reads was performed with FastQC (Andrews, 2010) and the mean quality value across each base position in the read was obtained with MultiQC (Ewels at al., 2016). As outlined in the workflow for Trimitomics (Fig. 1), our pipeline is comprised of three sequential steps using different software that are used stepwise, depending on the success of mt genome assembly in the preceding step. The first step consists of the use of the NOVOPlasty version 2.7.1 organelle assembler (Dierckxsens et al., 2016), with a range of *k*-mer distributions, using the raw reads as input. If the full mt genome is not recovered, and no or only a partial mt genome is obtained, the raw reads are then cleaned using Trimmomatic 0.33 (Bolger et al., 2014) with the following settings: ILLUMINACLIP:./Adapters.fa:2:30:10 LEADING:3 TRAILING:3 SLIDINGWINDOW:4:20 MINLEN:30, with the Adapters.fa file adjusted to include the adapter sequences specific to each read pair. RNA-Seq reads are then mapped to their respective “reference” genome (the closest mt genome published) with Bowtie2 (Langmead et al., 2012), using default settings. Mapped reads are then assembled *de novo* with genome guided Trinity version 2013_08_14 (Grabherr et al., 2011; Haas et al., 2013).

**Figure 1.**
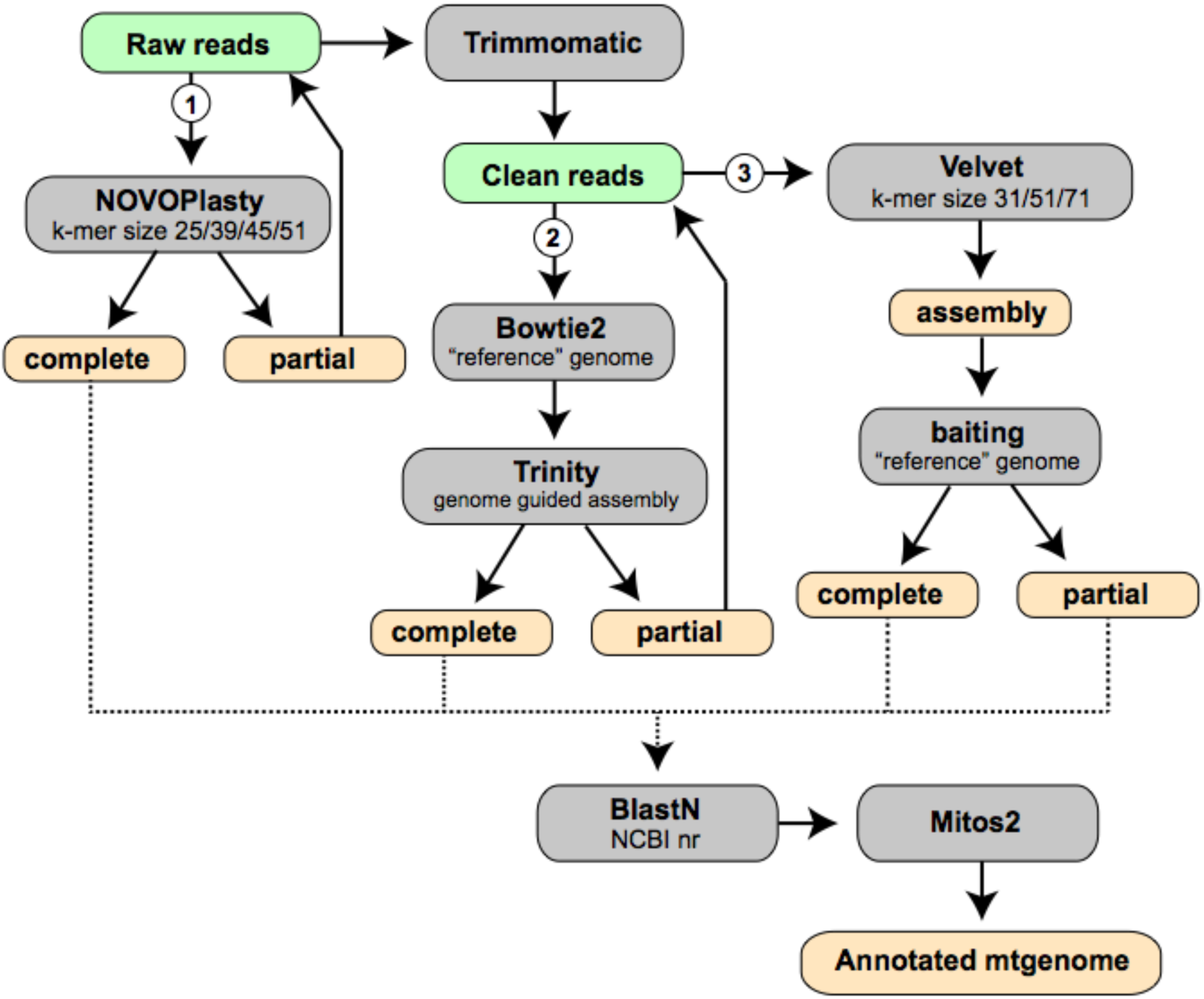
Proposed Trimitomics pipeline workflow. NOVOPlasty organelle assembler with range of k-mer distributions is applied on raw reads (1). Preferably a complete circularized mt genome is acquired or several contigs are obtained. In the latter case, resulting contigs, are merged into supercontigs by mapping to the reference mt genome to recover the complete mt genome. If mt genome is still partial we quality trim the raw reads with Trimmomatic and use them in reference-guided Bowtie2 alignment (2). Thereafter, mapped reads corresponding to the mt genome are extracted and further assembled with genome guided Trinity assembly. If mt genome is still partial we use velvet assembly (3) with range of k-mer distributions. From resulting assembly, mt genome contigs are extracted baiting with reference mt genome. If possible, resulted contigs are assembled into supercontigs. If none of the proposed methods acquired complete mt genome all resulting contigs were merged into supercontigs in order to get best results. In the final steps, supercontigs are checked for homology with BlastN in NCBI nr database and annotated using Mitos2.

If neither of the above-mentioned methods successfully retrieved mt genomes, then the whole transcriptome is assembled using Velvet 1.2.10 (Zerbino & Birney 2008) with a range of k-mer sizes (31,51,71). Mitochondrial contigs were then mined from *de novo* transcriptome assemblies with BlastN (Altschul et al., 1990) using the “reference” genome for the analysed species.

If the mt genome is not recovered by any of the three methods, the results are combined as a meta-assembly in order to obtain the best results. When the retrieved mt genomes are obtained in several contigs, this meta-assembly could be performed in Geneious v. 10.2.4, but can be done manually by comparison of contig ends using any alignment software. When the complete or almost complete mt genome is obtained, the assembly data is checked for homology with BlastN against the NCBI nr database. Further annotation is then performed on the web server MITOS2 (Bernt et al., 2013) using the appropriate translation table.

In the cases of *C. candelabrum, B. glabrata* and *P. dumerilii* the assembled and annotated mt genomes acquired with our pipeline were aligned with their respective published genomes using MAFFT (Katoh et al., 2002) to estimate the accuracy of the mt genome retrieval.

For all assembled mt genomes, data alignment statistics were obtained with SAMTOOLS (Li et al., 2009) after the RNA-seq reads were mapped to their assembled mt genome with Bowtie2. Potential PCR duplicates were marked and subsequently removed with the Picard tool (http://broadinstitute.github.io/picard). Uniquely mapped mitochondrial reads in the transcriptomes were determined with the RSeQC package with a minimum mapping quality 30 (Wang et al., 2012). Alignment statistics were plotted using the ggplot R package in Rstudio (Wickham, 2016). Coverage graphs were obtained in Ugene (Okonechnikov et al., 2012).

We computed linear regressions between the number of reads and the percentage of mt genome recovery for each of the methods to understand whether there existed a relationship between those variables, using the R package *lattice* (CRAN.R-project.org, Sarkar, 2008).

### Results

#### Quality of RNA-seq data

The quality control checks on raw sequence data for all analyzed transcriptomes are summarized in Figure 2. Most reads of all datasets showed average quality scores (Phred score) in the range 30–40, with the sponge *C. candelabrum* as the sole exception. In this species, a Phred score plateau of 20 was observed at the beginning and at the end of the reads (Figure 2). The majority of the analyzed raw reads were up to 100 bp in length, except for *S. officinalis* and *A. valida*, which were up to 150 bp (Figure 2).

**Figure 2.**
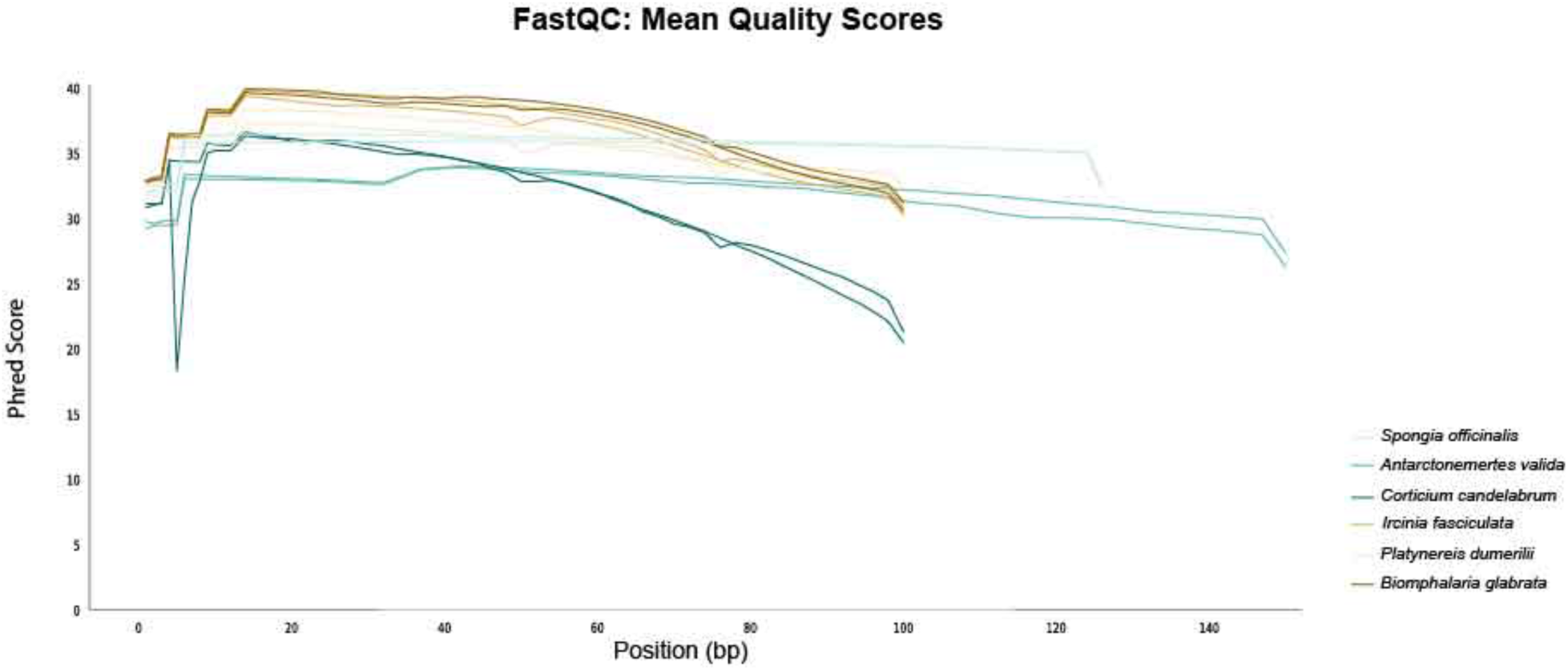
Quality assessment of all analysed raw reads expressed as mean quality value across each base position in the read. Each species, labelled with different colour as outlined in the legend, has two data sets for forward and reverse paired reads.

#### Assembling mitochondrial genomes from RNA-seq data

In this study, we aimed to retrieve mt genomes from transcriptomic data comprising four phyla, Porifera, Nemertea, Mollusca and Annelida. Efficiency of the proposed Trimitomics workflow is summarized in Table 1 and discussed in detail below. It is important to note that there was no correlation between the number of input reads and the percentage of recovery of the mt genome for any of the three steps of Trimitomics (Supplementary File 1).

**Table 1.** Summary of efficiency of the proposed workflow. Newly obtained mt genomes are marked in **bold font**. Asterisk (*) is denoting that genome size is estimated on the median size of published mt genomes from closely related species.

| PHYLUM | SPECIES | RAW READS IN<br>RNA-SEQ DATA<br>(paired end) | MT-READS<br>COUNT IN RNA-SEQ DATA (%) | MT GENOME RECOVERY<br>(%)<br>NOVOPlasty-Bowtie2/Trinity-Velvet | ACCURACY (%)<br>NOVOPlasty-Bowtie2/Trinity-Velvet |
| --- | --- | --- | --- | --- | --- |
| PORIFERA | *Corticium candelabrum* | 35,880,718 | 0.28 | 69.4/66.8/57.0 | 90.6/92.8/94.5 |
|  | ***Spongia officinalis*** | 237,867,408 | 0.04 | 100/79.0/96.0* | New mt genome |
|  | ***Ircinia fasciculata*** | 43,587,867 | 0.43 | 72.0/41.0/87.6* | New mt genome |
| NEMERTEA | ***Antarctonemertes***<br>***valida*** | 113,745,688 | 0.28 | 20.9/82.0/59.0* | New mt genome |
| MOLLUSCA | *Biomphalaria glabrata* | 52,648,074 | 2.94 | 97.0/91.0/53.0 | 99.6-99-84.6 |
| ANNELIDA | *Platynereis dumerilii* | 78,789,842 | 27 | 10.0/76.7/21.5 | 99.3/94.9/89.7 |

#### Porifera

For *C. candelabrum* NOVOPlasty with k-mer size 39 resulted in one contig of 12,773 bp in size, thus recovering 69.4% of the expected genome size. This assembly of the mt genome showed 90.6% sequence similarity with that published previously. When using the Bowtie2/Trinity combination, we obtained 8 contigs with mitochondrial similarity (size: 215–6,335 bp). The final meta-assembly of these contigs in Geneious allowed us to recover 12,297 bp (66.8% of the total genome size) in two contigs. The similarity percentage with the published mt genome in this case was 92.8%. Finally, our Velvet assembly with *k*-mer size 71 initially gave 10 contigs (size: 339–4,293 bp), that were later assembled in a contig that recovered 57% of the mt genome with 94.5% accuracy. When we combined the results from NOVOPlasty and Bowtie2/Trinity, we obtained an almost complete mitochondrial genome (99.4% mt genome recovery) with only 120 bp missing within an intergenic region (Fig. 3).

**Figure 3.**
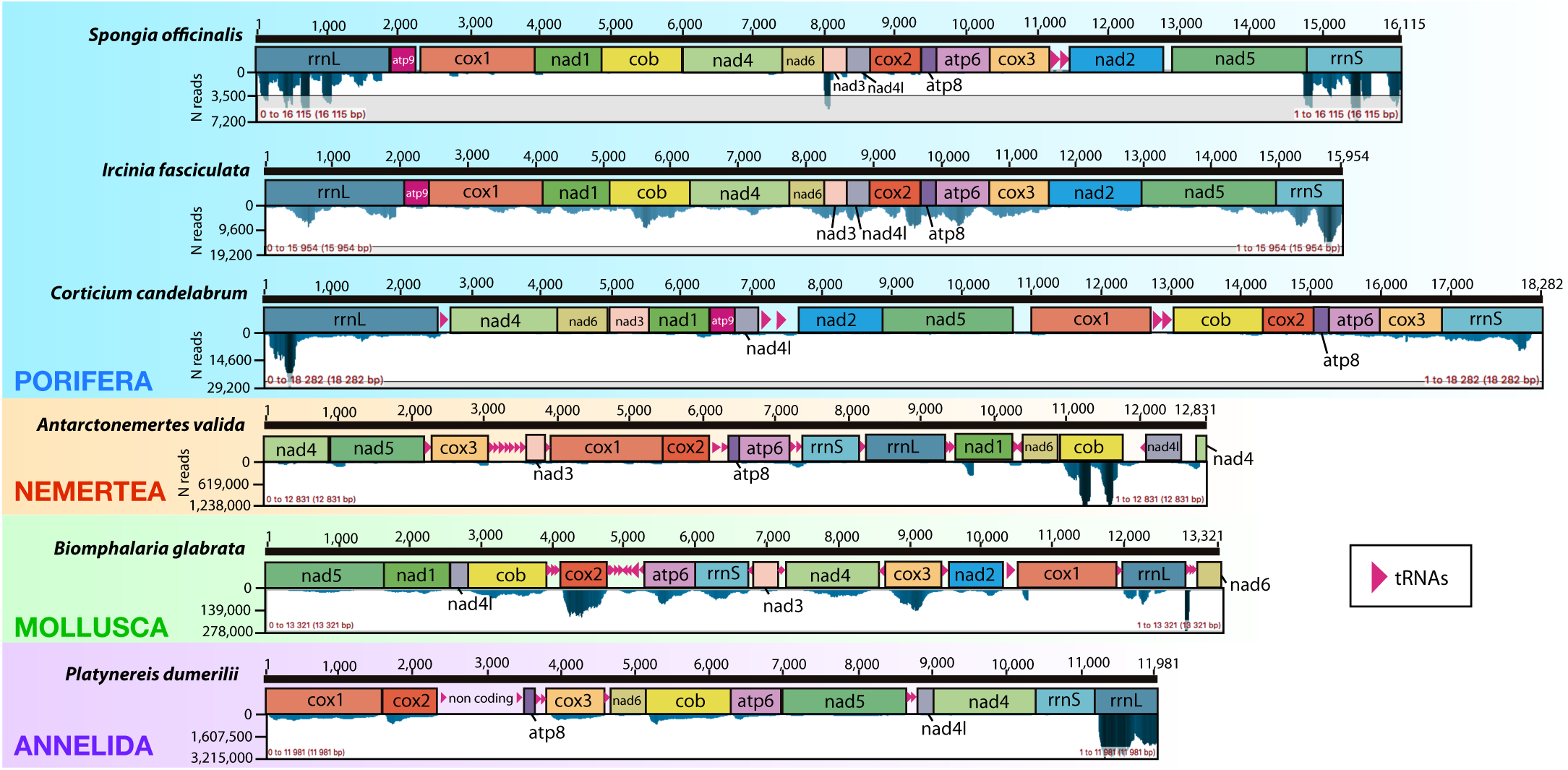
Gene order and read coverage for the obtained mt genomes from the six species analysed in this study. Upper bar represents the size of retrieved mt genomes. Below is annotation with genes, rRNAs and tRNAs. Mitochondrial expression patterns are shown as coverage graphs below.

In the case of *S. officinalis*, the complete mt genome was recovered by NOVOPlasty alone (Fig. 3). The circularized 16,115 bp long mt genome is the first representative published for the genus *Spongia*. The mt genome of the closest related species *Hippospongia lachne* (family Spongiidae) was used as a reference, and the MAFFT alignment of both species, *S. officinalis* and *H. lachne*, showed 57% identity. The Bowtie2/Trinity combination initially resulted in 21 contigs (size: 201–8,649 bp). The meta-assembly of such contigs in Geneious gave 12,726 bp in 2 contigs. Finally, Velvet using a *k*-mer size of 71 gave 26 contigs with lengths 71–1,893 bp, that were further assembled into 3 contigs (13,945 bp), which represented 96% of the mt genome (Table 1).

For *I. fasciculata* the expected mt genome size was roughly estimated from the median size of already published mt genomes from the congeneric *Ircinia strobilina* (NC_013662.1) and *Ircinia* sp. (KC510273.1) to be approximately 16,000 bp. NOVOPlasty (*k*-mer size 45) assembly resulted in 3 contigs with length ranging from 314–9,639 bp. The meta-assembly of these 3 contigs resulted in one contig of 11,669 bp. The Bowtie2/Trinity combination using *Ircinia strobilina* as reference mt genome gave 11 contigs that could be assembled into 3 contigs with 6,642 bp in total. Velvet (*k*-mer 71) gave 14 contigs (size: 225–5,553 bp) that were assembled for the final mt genome of 13,945 bp in 3 contigs. When the results from NOVOPlasty and Velvet were combined we attained 98.3% of mt genome recovery with a sequence length of 15,954 bp and all protein coding genes recovered (*nad2* partial at the 5’ end), as well as rrnS and rrnL. Some parts of one intergenic region and 2 tRNAs were missing (Fig. 3).

#### Nemertea

For *Antarctonemertes valida*, the closest related species with already published mt genome is *Gononemertes parasita* from the same suborder *Eumonostilifera*. Thus, this nemertean species, with mt genome size of 14,742 bp was used as reference genome.

With NOVOPlasty at a range of *k-*mer sizes only 3,088 bp were obtained. With Bowtie2/Trinity initially gave 27 contigs of sizes 213–2,440 bp. Velvet *de novo* assembly, using all 3 *k*-mer sizes, a total of 8,884 bp in 12 contigs were recovered. Final meta-assembly recovered a partial mt genome with 12,298 bp in 10 contigs representing 82% of the expected mt genome, with only *nad2* completely missing. In addition, *nad1, nad4, nad5* and *cox2* were partially obtained and 9 tRNAs were missing.

#### Mollusca

For *B. glabrata*, 97% of the genome was recovered from the raw sequencing reads with NOVOPlasty (k-mer size 39) in one contig, with accuracy 99.6% respect to the published mt genome, leaving only 616 bp unaccounted for. Bowtie2/Trinity resulted in 5 contigs with length ranging from 447–5,195 bp, thus recovering 91% of the mt genome. The meta-assembly of these 5 contigs resulted in 12,447 bp in one contig with 99% accuracy. Finally, *de novo* assembly from Velvet gave 19 contigs with length from 141–1,329 bp with 84.6% accuracy. For final analysis, we created a meta-assembly with the results obtained from NOVOPlasty and Velvet, which resulted in 97.5% mt genome recovery with 91.1% accuracy. All protein coding genes were recovered, with a small 5’ partial deletion from *cox1* and 3’ partial deletion of *nad6,* and 9 tRNAs missing out of 22.

#### Annelida

For *P. dumerilii,* NOVOPlasty assemblies with *k*-mer size 25, 39 and 45 gave poor results (<600bp), although *k*-mer size 51 gave results of 1,573 bp in one contig with 99.3% accuracy. Velvet assembly resulted in 3,353 bp in 23 contigs (with size ranging from 141–251 bp) with 89.7% accuracy. The combination of Bowtie2/Trinity resulted in 17 contigs (size: 273–2528 bp). Final meta-assembly combining all of the above data sources resulted in 10 contigs encompassing 76.7% of the genome with 94.9% accuracy. Sequences for *nad1, 2* and *3* could not be recovered, and neither were the majority of tRNAs (recovered only 6 out of 22 tRNAs). We were surprised at the poor assembly of the *P. dumerilii* mt genome, given the high percentage of reads with mitochondrial identity (Table 1), but these could represent reads from a diverse range of individuals, or alternatively could represent contamination from other eukaryote sources, which would both impair our assembly.

#### Read mapping and coverage for the mt genome

Our analyzed transcriptomic data contained diverse percentage of mitochondrial-derived reads (Table 1). The majority of the datasets had less than 1% of mt genome reads represented in the overall sequencing reads (Table 1). Interestingly, even though the mt genome of *Platynereis dumerilii* was the least complete of our final assemblies, a large percentage of the total number of reads (27%) was identified as being of mitochondrial origin (Table 1). The alignment statistics of exclusively mitochondrial reads are presented in Figure 4. Percentage of duplicate reads in the studied RNA-seq datasets was around 7% for most species (Fig. 4), with the exception of *Spongia officinalis* and *Corticium candelabrum* (up to 40 and 24%, respectively). The number of removed reads and those uniquely paired to the analyzed mt genome was quite similar among the different taxa used in our study (Fig. 4).

**Figure 4.**
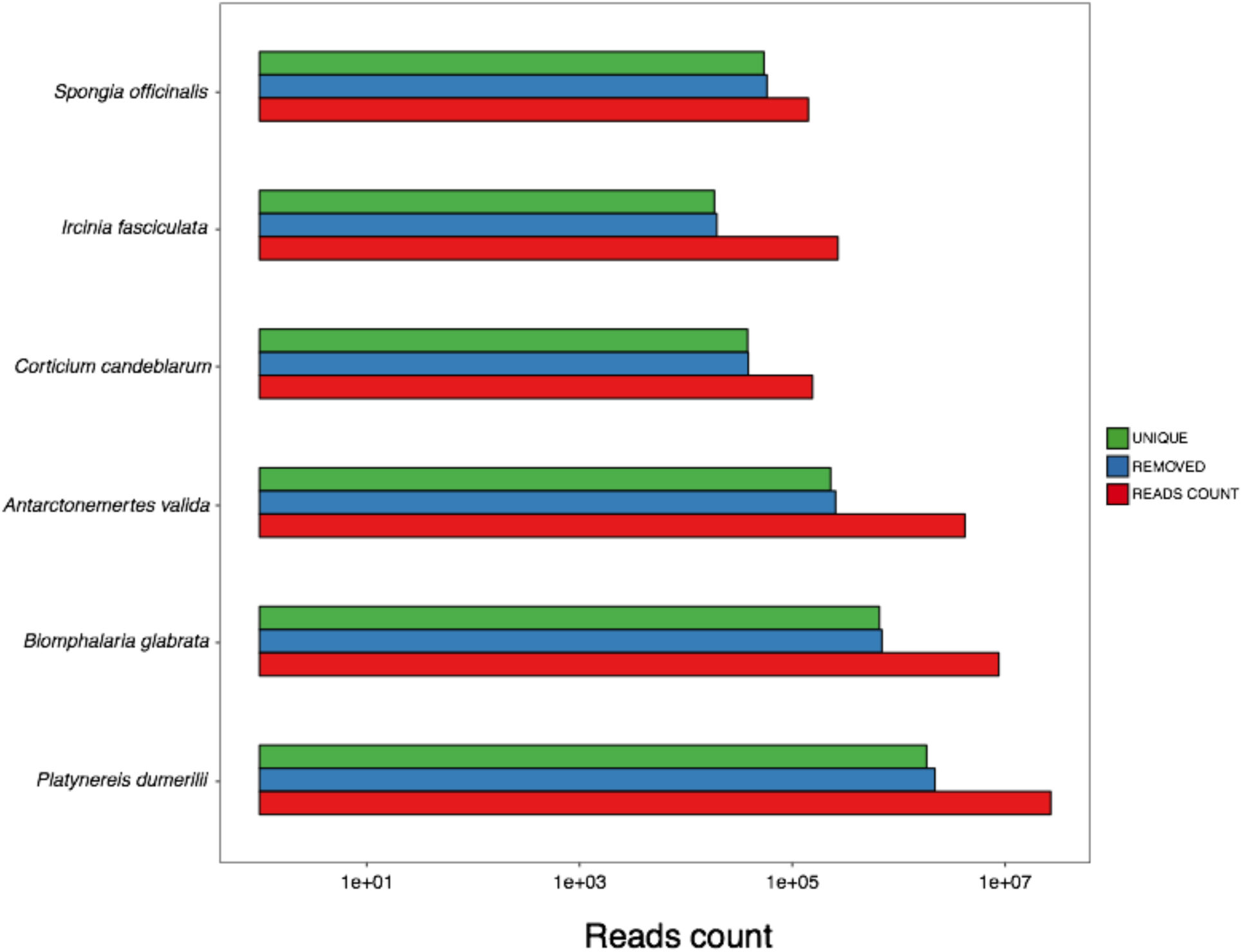
Alignment statistics for the number of reads obtained for the six species analysed in this study. Total mitochondrial reads count in analysed transcriptomes (reads count, in red). After filtering potential PCR duplicates (removed, in blue) and those that are uniquely mapped (unique, in green).

It is not quite clear whether these reads are artefactual duplicates that occurred during library construction or indeed represent independent mitochondrial reads in the transcriptomic data. Mt genomes are relatively small and are present in high copy number in cells, with limited numbers of potential annealing sites, and these percentages could reflect current situation in the analyzed species. Nevertheless, to ensure unbiased analysis for further studies these reads were removed.

Mapping of the RNA-Seq data onto the retrieved mt genomes gave a first insight into the transcription of the mitochondrial genes in our target invertebrates. In most cases both small and large ribosomal genes (*rrnS* and *rrnL*) were more expressed than the protein coding genes, although the number of reads mapped to those ribosomal genes varied substantially among datasets (Table 2 and Fig. 3). Only in the nemertean *A. valida,* a protein coding gene *cytochrome oxidase b* (*cob*) was more expressed than ribosomal genes (Table 2 and Fig. 3). In the sponges *S. officinalis* and *I. fasciculata*, the protein coding genes *nad3, nad4L*, and *atp8*, were the next-most covered by RNAseq data (Table 2 and Fig. 3), whereas in *C. candelabrum* the most represented was *cytochrome c oxidase 3* (*cox3*), albeit with very low expression values (Table 2 and Fig. 3). In *B. glabrata* the next most-expressed mitochondrial genes (after rRNA) were *cytochrome c oxidase subunit 2* and *3* (*cox2* and *cox3*), followed by *cob* and *atp6*. In the annelid *P. dumerilii,* after the highest expression values exhibited by the large ribosomal gene (*rrnL*), the cytochrome genes were the next most represented portion of the assembly (Table 2 and Fig. 3.). These values can be used to approximate expression, but should not be taken as definitive proof of expression values due to the poly-A selection steps incorporated into library construction, which will favour transcripts proximal to polyadenylation signals, whether incorporated by RNA polymerases or natively present within the raw mitochondrial sequence.

**Table 2.** Number of reads aligned to each of the genes in the mitochondrial genomes.

| Species | atp6 | atp8 | atp9 | cob | cox1 | cox2 | cox3 | nad1 | nad2 | nad3 | nad4 | nad4l | nad5 | nad6 | rrnL | rrnS | TOTAL |
| --- | --- | --- | --- | --- | --- | --- | --- | --- | --- | --- | --- | --- | --- | --- | --- | --- | --- |
| *Spongia officinalis* | 225 | 129 | 94 | 326 | 606 | 233 | 211 | 261 | 300 | 302 | 444 | 202 | 564 | 201 | 999 | 600 | 5,697 |
| *Ircinia fasciculata* | 1,963 | 792 | 202 | 2,878 | 1,526 | 1,710 | 1,843 | 1,368 | 790 | 1,223 | 1,437 | 915 | 1,799 | 906 | 2,897 | 3,225 | 25,474 |
| *Corticium candeblarum* | 2886 | 614 | 714 | 2088 | 1950 | 1421 | 5054 | 2035 | 1915 | 379 | 985 | 1176 | 1561 | 568 | 17918 | 11762 | 53,026 |
| *Antarctonemertes valida* | 25,029 | 2,260 | - | 76,021 | 42,911 | 7,540 | 19,548 | 15,231 | 8,694 | 65 | 23,640 | 5,259 | 39,642 | 13,388 | 15,200 | 20,201 | 314,629 |
| *Biomphalaria glabrata* | 57,634 | 5,061 | - | 73,889 | 3,646 | 84,668 | 73,662 | 34,870 | 21,100 | 4,914 | 60,840 | 6,263 | 37,909 | 1,134 | 44,561 | 46,411 | 556,562 |
| *Platynereis dumerilii* | 121,189 | 3,771 | - | 360,465 | 533,632 | 188,164 | 197,631 | - | - | - | 197,010 | 41,758 | 314,525 | 41,059 | 509,018 | 48,474 | 2,556,696 |

## Discussion

The ‘omics’ revolution and the development of novel bioinformatic tools that are now routinely available at little to no cost to all laboratories has provided a wealth of data from almost all animal groups. Nowadays, genome skimming and RNA-seq projects are feasible for non-model organisms and the SRA database grows larger on a daily basis. Despite some awareness that RNA-seq data can contain high numbers of mt genome reads (Smith, 2013), the majority of available methods for mt genome assembly are customized for recovering these from genome sequencing projects (either whole genome sequencing or genome skimming). But recently, the mining of mt genomes from RNA-seq data has been successfully performed in a handful of eukaryotes, including algae (Tian & Smith, 2016), reptiles (Lyra et al., 2017) and sea urchins (Dilly et al., 2015), yet using disparate software approaches, including Bowtie2/Trinity (Tian & Smith, 2016), MIRA v4.0/MITObim v1.8 (Lyra et al., 2017), and blast against the mt genomes of closely related species (Dilly et al., 2015). Here we propose a standardized workflow for recovering at least the coding regions of mt genomes from RNA-seq data across Metazoa.

Using data from a number of non-model invertebrates we recovered with our pipeline more than 97% of the mt genome in the majority of cases (Fig. 3). In our proposed Trimitomics workflow (Fig. 1) NOVOPlasty is the first tool implemented, and was empirically superior in runtime, memory consumption and accuracy when compared with other existing tools for mitochondrial assembly. This is expected, as it is a newly published pipeline which incorporates advancements in bioinformatics tools targeting the mt genome specifically (Dierckxsens et al., 2016).

The Bowtie2/Trinity approach has been used previously to recover mt genomes (Tian & Smith, 2016), adding the coverage and expression levels of each of the genes in the mitochondrial chromosomes (Tian & Smith, 2016), and it has proven useful in this analysis as well. For the nemertean *A. valida* and the annelid *P. dumerilii*, Bowtie2/Trinity yielded the best results. However, the initial identification of reads as being of mitochondrial origin depends on inference of homology using Bowtie2 mapping, and thus requires a relatively closely-related sequence to act as “bait”. Without this inference of homology, reads are simply excluded from the following Trinity assembly. We therefore recommend that if “unspannable” gaps are found when using the Bowtie2/Trinity approach, a *de novo* assembly (either with Trinity or Velvet, but using all reads) is assayed and contigs from these are tested to see if they can be used to span such problematic areas.

Velvet is the most memory and time consuming approach, and thus we recommend that it is performed after previous approaches have been trialled, but in many cases it yields significant results. For example, in the case of *I. fasciculata* Velvet increased the final mt genome assembly by 27%. As Velvet is an assembler that was designed primarily for genomic read data, it is optimised to assemble reads of relatively uniform coverage. It can therefore be optimised further for assembly of specific regions of the mitochondrion using the –min_cov, -max_cov and –exp_cov settings once general coverage levels are known, but this can be a time-consuming approach, and is beyond the scope of this study.

Efficiency in the recovery of the complete mt genomes, as expected, depends on RNA-seq library preparation, quality of read data and the depth of sequencing. In this paper, it was not our goal to compare the suitability of available methods of library preparation, data quality or sequencing depth in the recovery of the mt genome but rather to demonstrate that the complete mt genome of understudied species could be obtained with our pipeline using a diverse set of RNA-seq data. The recovery of complete (rather than CDS-containing only), mt genomes from RNA-seq datasets could be due to either genomic contamination, incomplete DNA digestion during RNA library preparation or could be an indication that the majority of the mt genome is actually transcribed (Tian & Smith, 2016), even if only transiently before RNA editing. Even the intergenic regions were recovered in RNA-seq datasets, which enabled complete mt genome recovery.

Thus, given that in many different laboratories RNA-seq experiments are performed more and more routinely, our Trimitomics pipeline might become a powerful approach to exploit this neglected goldmine of novel mitochondrial genomes within RNA-seq data and ensure their future usage to address various biological and evolutionary questions. In any case, the recovery of even partial mt genomes with our pipeline in non-model invertebrates represents a significant contribution, as in these taxa mitochondrial genome data is scarce. We have shown that even when the mitochondrial read percentage is low, such as 0.04% in case of *S. officinalis,* complete mt genomes can be obtained (Table 1).

mRNA read data generated using standard methods are not sufficient to fully characterize mitochondrial expression patterns, as most library preparation methods target transcripts with a poly-A tail, which is not universally the case in mitochondrial genes. However, it can provide information regarding general transcriptional trends, especially on a within-species basis, that can be explored further. Little has been published about the baseline gene expression levels of mitochondrial genes in invertebrates (e.g., Wang et al., 2013; Perera et al., 2016).

As expected, and previously reported for other eukaryotic groups (e.g., Tian & Smith 2016; Wang et al., 2013; Perera et al., 2016), for previously analyzed mt genomes, coverage in our datasets was lowest in intergenic regions and highest in areas encoding rRNAs (Fig. 3). Interestingly, our results for the transcriptional landscape across invertebrate phyla indicates that the transcriptional profiles are diverse, even among the three analyzed poriferan species. Besides the ribosomal genes (especially *rrnL*), the cytochromes (*cox1, cox2,* and *cox3*) and *atp6* are usually the most transcribed in animal groups (e.g., Wang et al., 2013; Perera et al., 2016). This was true for the sponge *C. candelabrum* (although only *cox3* and *atp6*), the mollusc *B. glabrata* and the annelid *P. dumerilii* (Fig. 3), but the two dictyoceratid sponges *S. officinalis* and *I. fasciculata* showed greater expression in the genes *nad3, nad4L, atp6* and *atp8* (Fig. 3). This indicates that mitochondrial genes are expressed according to the animal species sampled, and, presumably, the current energy requirements of the sample taken.

By way of example, although ribosomal genes are usually the most expressed genes within the mitochondria in the lepidopteran *Helicoverpa zea*, in its embryos *cox1* is by far the most highly expressed gene (Perera et al., 2016), while in the vole *Clethrionomys glareolus cox1, cox2, cox3* and *atp6* are expressed more than ribosomal genes (Markova et al., 2015). Our pathway provides a means to determine this information in a more systematic manner in any previously published RNA-seq experiment, provided library construction was performed in an internally-consistent manner, and thus provides a pathway to understanding the mitochondrial transcriptional landscape in any interesting biological framework, either by leveraging extant resources, or through novel investigations in the future.

## Conclusions

Mitochondrial data are of diverse utility, and can be particularly crucial for investigations into the biology and evolution of non-model organisms. Here we have demonstrated a means of rapid and relatively inexpensive derivation of coding and, occasionally, full length mitochondrial sequences from a range of non-model species. This method will allow new information to be leveraged from extant datasets, with minimal or no cost, which will open doors for investigations in even the most recalcitrant of clades.

## Acknowledgements

The authors thank Dr Cristina Diez for her helpful discussion and support and Dr Peter Foster for IT support. This work was supported by a grant from the Villum Foundation (grant number 9278), NJK was supported by a H2020 MSCA grant during manuscript preparation [IF750937].

## Data Accessibility Statement

Raw reads for *Antarctonemertes valida* and *Spongia officinalis* were deposited at SRA with accession numbers SRP157324 and SRP150632. The newly assembled mt genomes *Spongia officinalis, Antarctonemertes valida* and *Ircinia fasciculata* were deposited at Genbank with the following accessions: MH768970-MH768972.

## Author Contributions

AR and BP conceived the study and gathered specimens and datasets. BP, ST, VK, MER and NJK designed and performed experiments. BP and MER performed assembly and bioinformatic analysis. AR provided the funding necessary to perform experiments and analyses. All authors wrote and agreed on the final form of the manuscript.

## Supporting/Supplemental Information

**Supplementary File 1.**
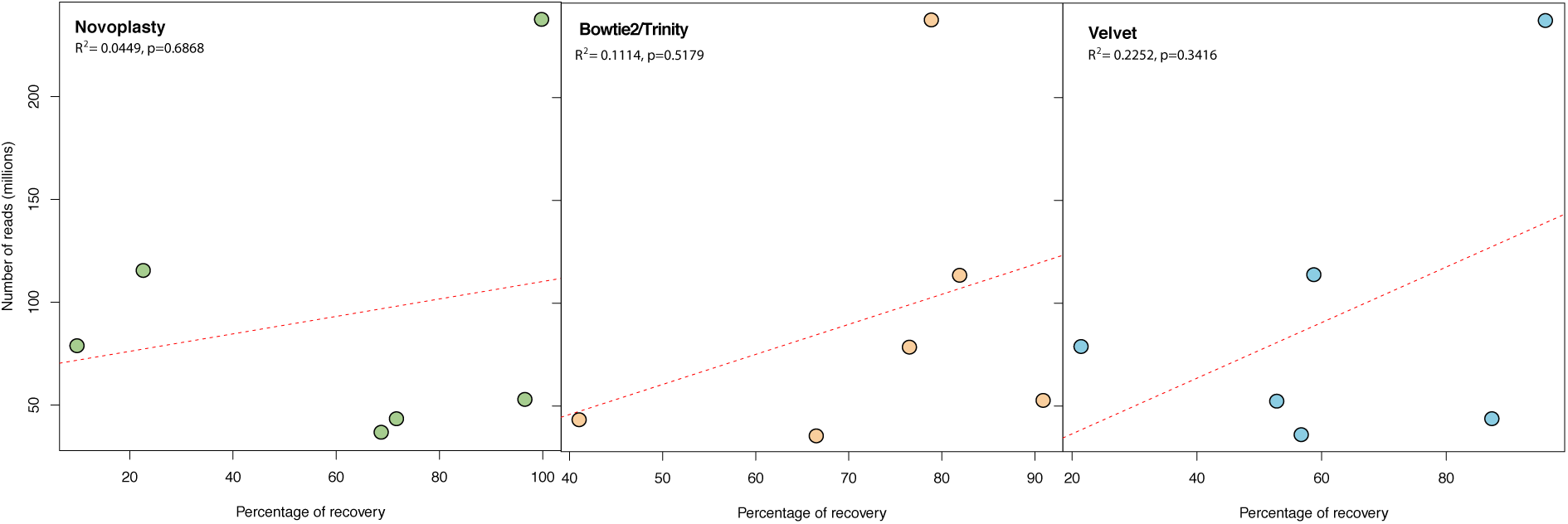
Correlation between number of reads used as input and the percentage of mt genome recovery for each of the three steps of Trimitomics.

